# *Egr1* loss-of-function promotes beige adipocyte differentiation and activation specifically in inguinal subcutaneous white adipose tissue

**DOI:** 10.1101/2020.06.03.131342

**Authors:** Marianne Bléher, Berbang Meshko, Rachel Gergondey, Yoann Kovacs, Delphine Duprez, Aurore L’Honoré, Emmanuelle Havis

## Abstract

Exercise, cold exposure and fasting lead to the differentiation of inducible-brown adipocytes, called beige adipocytes, within white adipose tissue and have beneficial effects on fat burning and metabolism, through heat production. This browning process is associated with an increased expression of the key thermogenic mitochondrial uncoupling protein 1, Ucp1. Egr1 transcription factor has been described as a regulator of white and beige differentiation programs, and Egr1 depletion is associated with a spontaneous increase of subcutaneous white adipose tissue browning, in absence of external stimulation. Here, we demonstrate that *Egr1* mutant mice exhibit a restrained *Ucp1* expression specifically increased in subcutaneous fat, resulting in a metabolic shift to a more brown-like, oxidative metabolism, which was not observed in other fat depots. In addition, *Egr1* is necessary and sufficient to promote white and alter beige adipocyte differentiation of mouse stem cells. These results suggest that modulation of *Egr1* expression could represent a promising therapeutic strategy to increase energy expenditure and to restrain obesity-associated metabolic disorders.

## Introduction

White adipose tissue (WAT) plays a crucial role in energetic homeostasis regulation through the storage of excess energy in a large central vacuole and also by performing metabolic functions and secreting several endocrine hormones such as Leptin (*Lep*) and Resistin (*Retn*) [1,2]. WAT is located in diverse sites throughout the body, such as the subcutaneous inguinal (SC-WAT), perigonadal (GAT), mesenteric (MAT) and perirenal (PAT) depots [3,4]. While these depots are collectively named white adipose tissue, they have different developmental origins: SC-WAT, GAT and MAT derive from lateral plate mesoderm in female mice whereas PAT derive from the somitic mesoderm [4]. These different origins could contribute to differences in the physiology of fat depots.

In contrast, brown adipose tissue (BAT) has a critical thermogenic function thanks to its property to dissipate excess energy through heat production [5]. BAT depots are derived from the somitic mesoderm [4] and are located in the inter-scapular region in mice and in both neck and supraclavicular regions in humans [5]. Brown adipocytes are smaller than white adipocytes. They exhibit multiple small lipid vacuoles and large number of mitochondria involved in heat production through the constitutive activation of the thermogenic uncoupling protein 1 (*Ucp1*) [6–8].

A third type of adipocyte, called beige adipocyte, has been described within WAT [5,9]. Morphological and transcriptomic analyses show that brown and beige adipocytes are remarkably similar and express the same thermogenic markers, including *Ucp1* [10]. *In vivo*, beige adipocytes differentiation occurs in response to long term cold exposition, fasting or exercise through a process referred to as browning [5,11]. White, beige and brown adipocytes have been shown to differentiate *in vitro* from adipose-derived mesenchymal stem/stromal cells (ASCs) isolated from lipoaspirates [12–15]. In this context, we previously demonstrated that mice lacking *Egr1* expression exhibit spontaneous browning of the inguinal subcutaneous white adipose tissue without external stimulation [16]. Egr1 is a multi-faceted transcription factor involved in the development and homeostasis of several organs [16–19]. Increased *Egr1* expression has been linked to obesity in both humans and murine models [18,19]. In contrast, *Egr1* knock-out mice display an increased energy expenditure and are protected from high fat diet-induced obesity and obesity-associated pathologies [19,20]. In addition, *Egr1* loss-of-function in mice induces spontaneous browning of inguinal SC-WAT, associated with *Ucp1* up-regulation [16].

In this study, we get further to first analyze the consequences of *Egr1* knock-out on *Ucp1* expression in all adipose tissue depots. Then, we tested whether *Ucp1* up-regulation in WAT comes with a shift to a BAT-like oxidative metabolism. Interestingly, we observed that *Ucp1* up-regulation in mutant *Egr1*^−/−^ adipose tissue was restricted to subcutaneous inguinal SC-WAT and associated with an increased oxygen consumption rate in this fat pad. To further characterize the role of *Egr1* in SC-WAT browning, adipose stem cells (mASC) were isolated from control *Egr1^+/+^* and mutant *Egr1^−/−^* SC-WAT and compared for their ability to differentiate into white or beige adipocytes *in vitro*. We observed that *Egr1* was necessary to promote white adipocyte differentiation and repress beige differentiation of mASC. *Egr1* gain-of-function experiments performed in C3H10T1/2 mesenchymal stem cells confirmed the positive role of *Egr1* on white adipocyte differentiation and its negative role on beige differentiation. Collectively, our results indicate that *Egr1* favors white adipocyte differentiation and represses SC-WAT browning. *Egr1* may thus become a promising target to combat obesity.

## Results and Discussion

### *Egr1^−/−^* mice display spontaneous increase of *Ucp1* expression specifically in SC-WAT

*Egr1* loss-of-function in postnatal and 4-month-old mice leads to a reduced body weight despite similar weight of SC-WAT fat pads in control *Egr1*^+/+^ and mutant *Egr1^−/−^* mice [16]. Body weight reduction was confirmed in 8-month-old *Egr1^−/−^* mice (Fig. 1A). In addition, all isolated white (subcutaneous, SC-WAT, gonadic, GAT, mesenteric, MAT and perirenal PAT) and brown (BAT) fat pads display similar percentage of body weight in control *Egr1*^+/+^ and mutant *Egr1^−/−^* mice (Fig. 1B, C). These observations indicate that lower body weight observed in mutant *Egr1^−/−^* mice is not due to a reduced adipose tissue mass, suggesting that *Egr1* loss-of-function reduces body weight through unknown global metabolic processes. *Egr1* loss-of-function has been shown to induce spontaneous browning of SC-WAT, associated with increased *Ucp1* expression, without external stimulation [16]. Here, we quantified the expression of *Egr1*, *Ucp1*, *Lep* and *Dcun1d3* in five different adipose tissue depots from control *Egr1*^+/+^ and *Egr1^−/−^* mice. As expected, *Egr1* was nearly undetectable in mutant *Egr1^−/−^* fat pads (Fig. 2A-E). A significant increase in *Ucp1* expression was observed in *Egr1^−/−^* SC-WAT compared to control (Fig. 2B). In contrast, *Ucp1* expression was not altered in the other 4 fat pads in mutant versus control (Fig 2A, C, D, E). These observations are consistent with previous results showing the absence of spontaneous *Ucp1* up-regulation in GAT from *Egr1*-deleted mice (Zhang et al., 2013). Our results indicate that *Egr1* deficiency induces a browning phenotype restricted to SC-WAT, and suggest that *Egr1* is required to repress SC-WAT browning in standard housing conditions. SC-WAT browning in mutant *Egr1^−/−^* mice is characterized by an increased number of beige adipocytes at the expense of white adipocytes [16]. *Leptin* (*Lep*) is a marker of white adipose tissue. This hormone regulates body weight, lipid and glucose metabolism through binding to its receptor which is expressed in the hypothalamus and several other tissues [21–23]. Consistently, *Lep* expression was reduced in mutant *Egr1^−/−^* SC-WAT (Fig. 2B). Taken together, our results suggest that Egr1 represses *Ucp1* and activates *Lep* expression in SC-WAT, which is in accordance with previous results showing recruitment of Egr1 to their promoters [16,24]. Decreased *Lep* expression in other WAT depots in *Egr1*-null mice (MAT, Fig. 2D) suggests that Egr1 can activate *Lep* independently of the browning phenotype.

**Figure 1.**
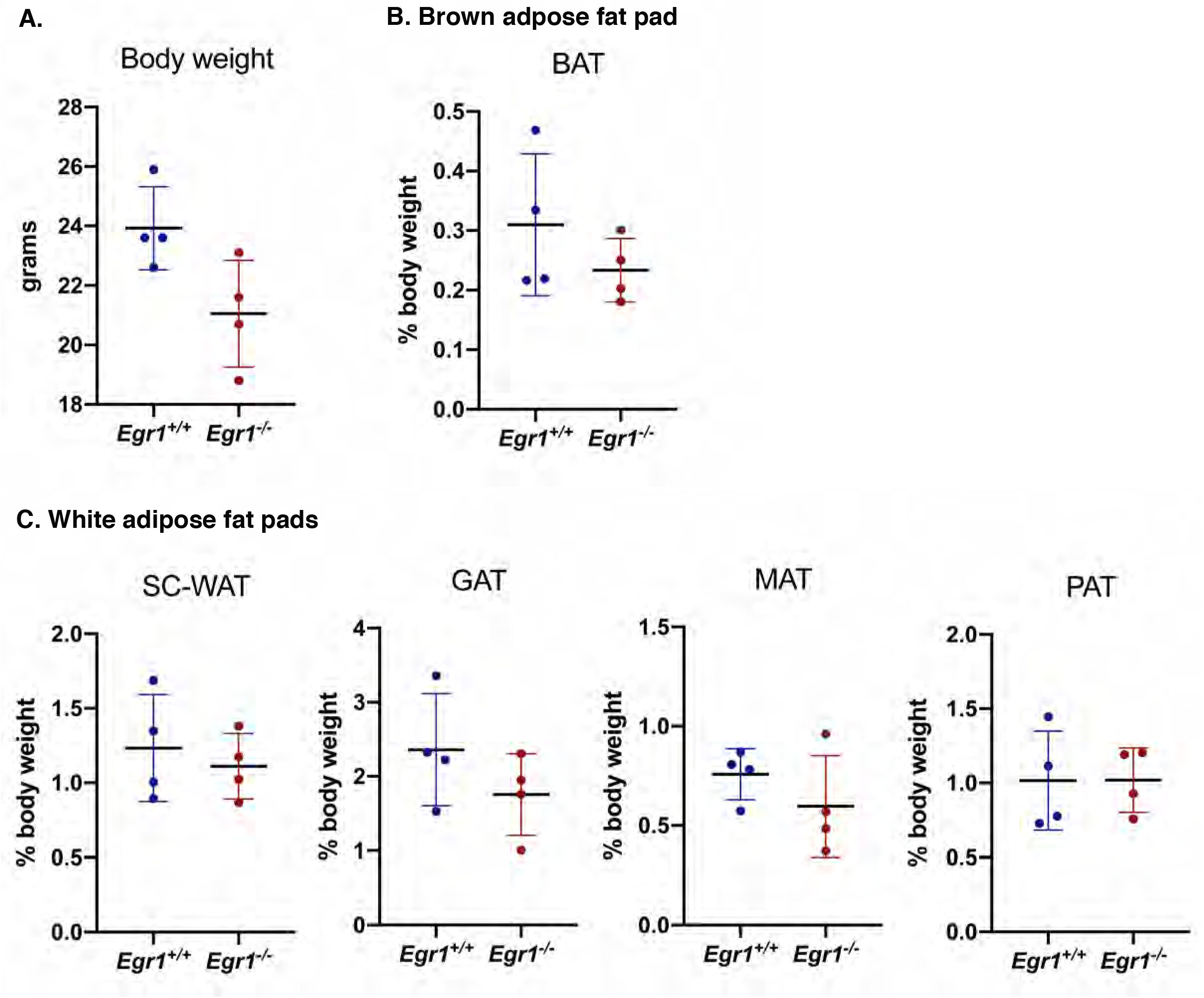
*Egr1* loss-of-function does not affect fat pad percentage of body weight. (**A**) Whole body, **(B)** Brown adipose tissue (BAT), **(C)** Inguinal sub-cutaneous adipose tissue (SC-WAT), gonadal adipose tissue (GAT), mesenteric adipose tissue (MAT) and perirenal adipose tissue (PAT) of 8-month-old *Egr1*^+/+^ and *Egr1^−/−^* female mice were weighted. Fat pad weights were represented as a percentage of body weight. Experiments were performed with n=4 for each genotype. Error bars represent the mean+ standard deviations, with *P<0.05.

**Figure 2.**
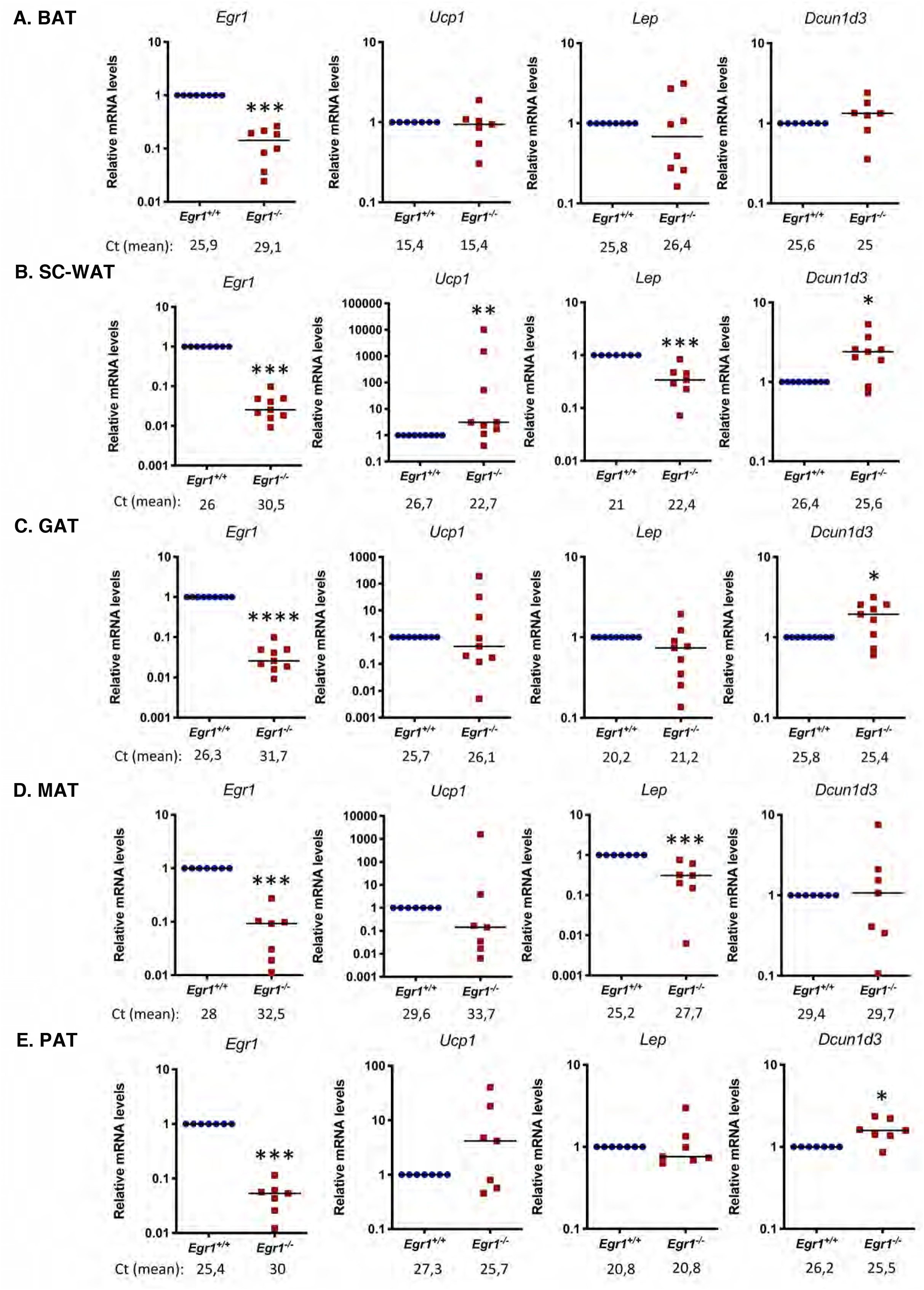
*Egr1* loss-of-function increases *Ucp1* expression specifically in SC-WAT. (**A**) BAT, **(B)** SC-WAT, **(C)** GAT, **(D)** MAT and **(E)** PAT of 8-month-old *Egr1*^+/+^ and *Egr1^−/−^* mice were used for RNA purification. RNA samples were analysed by RT-qPCR for *Egr1*, *Ucp1*, *Lep* and *Dcun1d3* expression. Experiments were performed with n=8 animals (BAT, MAT, PAT) or n=9 animals (SC-WAT and GAT) for each genotype. Transcripts are shown relative to the level of *18S* transcripts, with control *Egr1*^+/+^ being normalized to 1. The relative mRNA levels were calculated using the 2^^-ΔΔCt^ method. Error bars represent the means + standard deviations. The p-values were obtained using the Mann-Withney test, with * p<0.05, ** p<0.01, *** p<0.001

*Dcun1d3* has been shown upregulated in mutant *Egr1^−/−^* SC-WAT [16]. *Dcun1d3* is expressed in adipose tissue (Fig. 2-Supplement 1), however its role has not been elucidated yet. While *Dcun1d3* was present in all adipose depots in controls (Fig. 2A-E), its expression was upregulated in *Egr1*-deficient SC-WAT, GAT and PAT (Fig. 2B, C, E). This suggests that while *Dcun1d3* may be related to SC-WAT browning, *Egr1* can also regulate its expression independently of the browning process.

Taken together, these results indicate that *Egr1* loss-of-function increases both *Ucp1* and *Dcun1d3* expression and decreases *Lep* expression in SC-WAT, resulting in spontaneous browning in this specific adipose tissue depot. One hypothesis could be that Egr1 interacts with specific co-regulators in fat pads, leading to the regulation of different genetic programs in each depot. The co-factors allowing Egr1 to regulate the browning program might be specifically expressed in SC-WAT.

### *Egr1* loss-of-function induces a metabolic shift in white adipose tissue, leading to a BAT-like oxidative metabolism

We then investigated whether the brown transcriptomic signature observed in mutant *Egr1^−/−^* SC-WAT was correlated with metabolic modifications. While control *Egr1*^+/+^ SC-WAT is characterized by a low mitochondrial mass, marked by a low mitochondrial/nuclear DNA ratio, we observed an increased mitochondrial mass in mutant SC-WAT (Fig. 3A), with a ratio being much similar to that of BAT. Such increase is specific to this tissue and was not observed in any other tested fat tissues (Fig. 3-Supplement 1A). To examine whether this increased mitochondrial mass has functional metabolic consequences, we assessed oxygen consumption rate (OCR) of BAT, SC-WAT and GAT freshly dissected from control and mutant mice. Using Seahorse XFe24 analyzer, we showed that while *Egr1* loss-of-function does not affect GAT (Fig. 3-Supplement 1B, C) or BAT OCR (Figure 3B, C), it leads to an increase of all markers of oxidative capacity, including basal and maximal OCR, ATP-linked OCR and non-mitochondrial OCR. Importantly, all rates quantified in mutant *Egr1^−/−^* SC-WAT were close to that obtained with BAT, demonstrating a metabolic shift of *Egr1^−/−^* SC-WAT towards a BAT-like oxidative metabolism. As expected, this increase of oxidative capacity does not lead to altered ROS production, as indicated by similar carbonylation protein levels (Fig. 3-Supplement 2D), thus demonstrating that the browning process observed in *Egr1* mutant occurs in physiological conditions.

**Figure 3.**
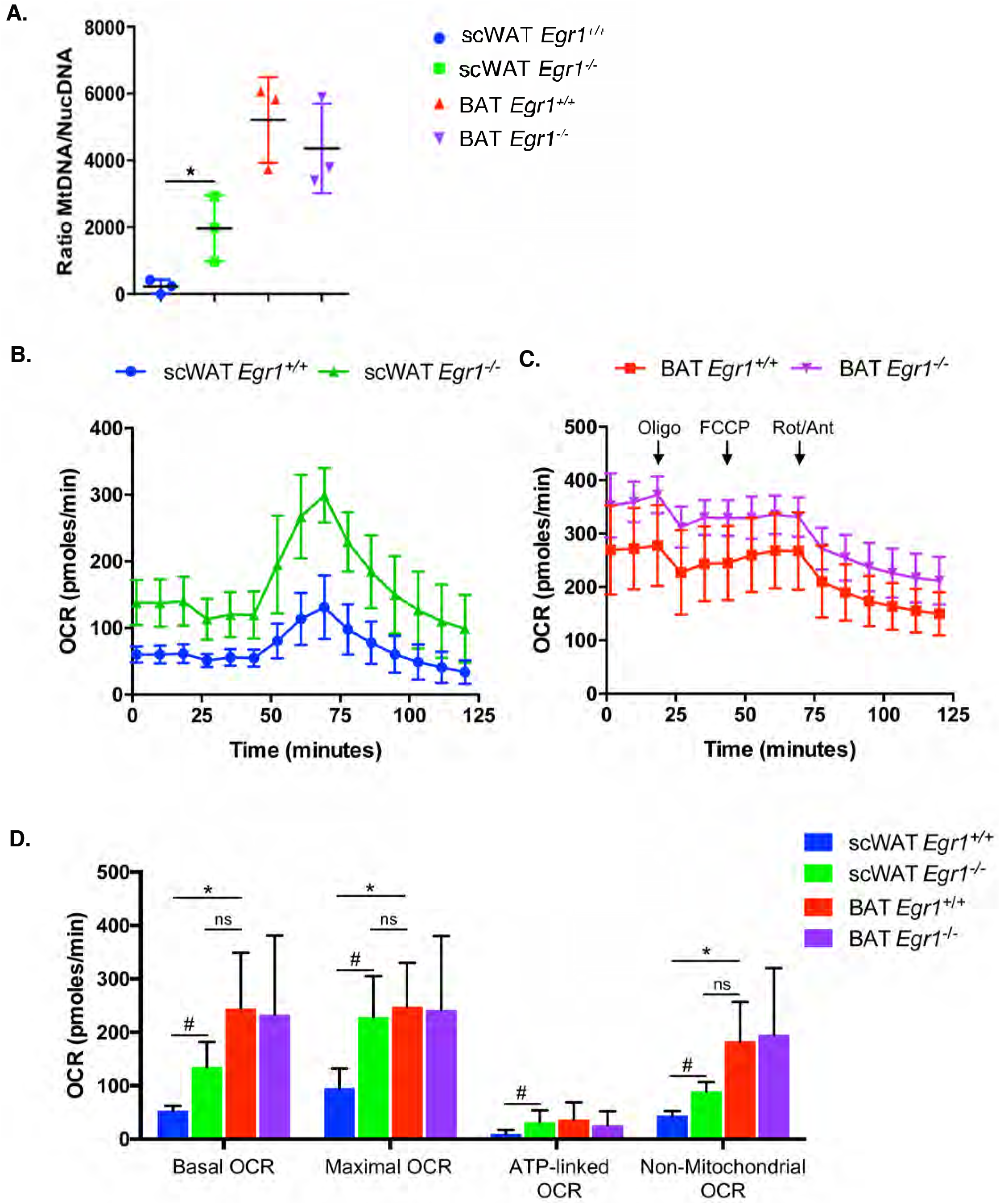
*Egr1* loss-of-function induces a metabolic shift in white adipose tissue, leading to a BAT-like oxidative metabolism. **(A)** WAT and BAT of 8-month-old control *Egr1*^+/+^ and mutant *Egr1^−/−^* female mice were dissected and used for DNA purification. Mitochondrial (*Cyt B*) and nuclear (*Ndufv1*) genes were quantified by qPCR and histogram represents their ratio. Error bars represent the means + standard deviations with n=3 animals for each genotype, *p<0.05. **(B, C)** Mitochondrial respiration measured by oxygen consumption rate (OCR) in basal conditions and after sequential addition of Oligomycin, FCCP, and a mix of Rotenone/Antimycin were simultaneously recorded on SC-WAT (B) and BAT (C) tissues, freshly dissected from 8-month-old control *Egr1*^+/+^ and mutant *Egr1^−/−^* female mice. **(D)** Histogram represents the basal OCR (determined as the difference between OCR before oligomycin and OCR after rotenone/antimycin A), maximal OCR (difference between OCR after FCCP and OCR after rotenone/antimycin A), ATP-linked OCR (difference between OCR before and after oligomycin), and the non-mitochondrial OCR (OCR after rotenone and antimycin A treatment) calculated from data obtained in A. Error bars represent the means + standard deviations with n=3 animals for each genotype, *p<0.05.

Our previous study has revealed a spontaneous browning of the SC-WAT in *Egr1* mutant, with *Ucp1* expression being up-regulated in absence of external stimulation [16]. We now show that this browning also exerts at physiological level with consequences on metabolic and bio-energetic activity of SC-WAT. Our results reveal a shift of *Egr1^−/−^* SC-WAT towards a BAT-like oxidative metabolism and are correlated with previous studies establishing a link between increased *Egr1* expression and obesity [18,19]. They suggest a central role of *Egr1* in regulation of SC-WAT homeostasis in the adult.

### *Egr1* is necessary to promote white and to repress beige adipocyte differentiation of mouse adipose stem cells

To further analyze the role of *Egr1* in spontaneous browning of SC-WAT in mutants, we tested whether *Egr1* loss-of-function can alter the differentiation of SC-WAT-derived mouse adipose stem cells (mASC) into white and beige adipocytes. mASC were isolated from inguinal SC-WAT of control *Egr1*^+/+^ and mutant *Egr1^−/−^* mice and subjected to white (Fig. 4A-B) or beige (Fig. 4C-D) adipogenic differentiation.

**Figure 4.**
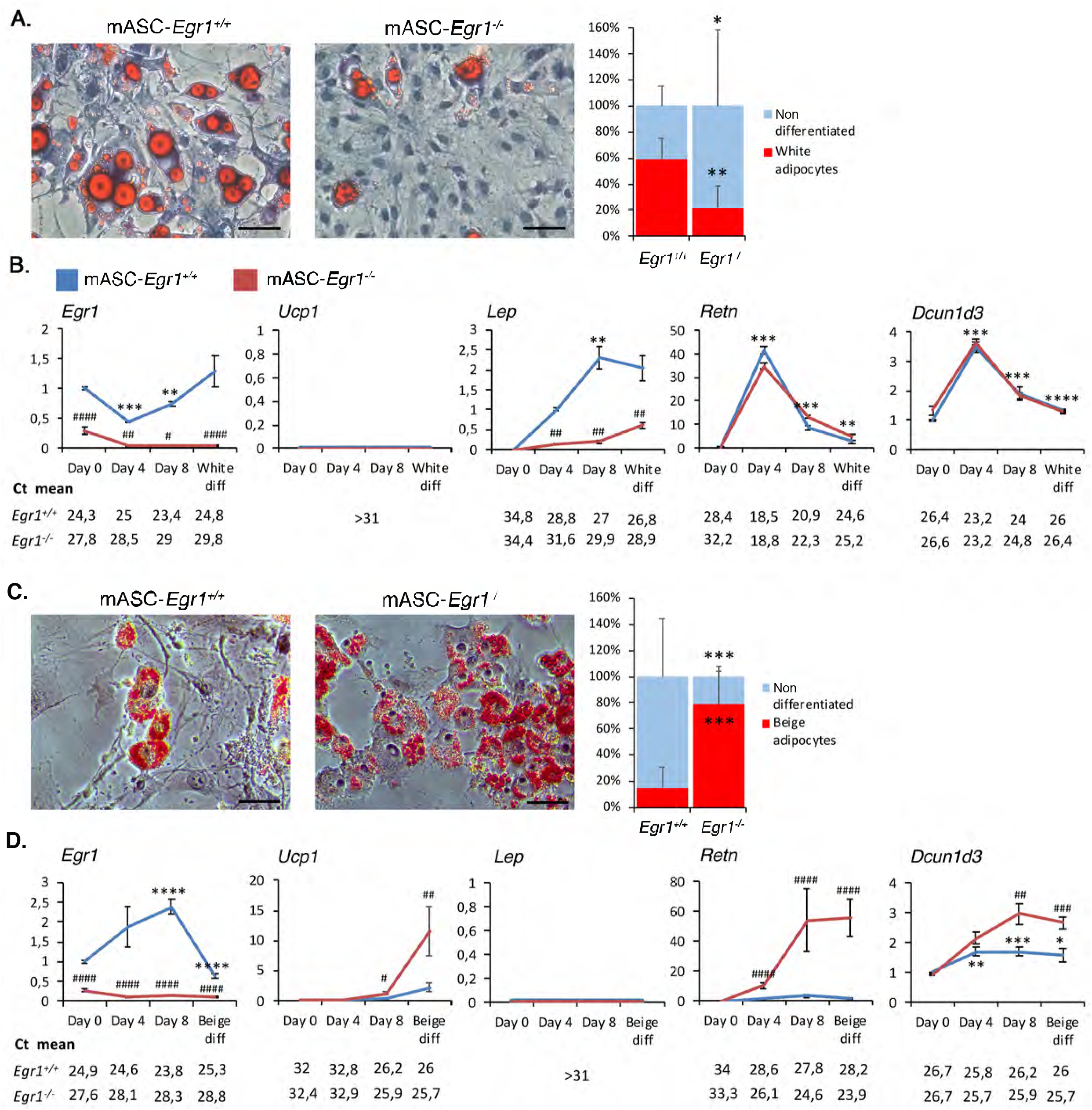
*Egr1* is necessary to promote white adipocyte differentiation and prevent beige differentiation of mouse adipose stem/stromal cells. **(A-F)** Control mASC-*Egr1*^+/+^ and mutant mASC-*Egr1^−/−^* were subjected to white (A,C,D) or beige (B,E,F) differentiation for 12 or 10 days, respectively. **(A, C)** Non-differentiated and white (A, right panel) or beige (C) adipocytes were stained with Oil Red O and Hematoxilin after white (A, left panel) or beige (C, left panel) differentiation and counted (A and C, right panels). For each genotype, results are expressed as a percentage of cells differentiated or not, counted from 7 pictures at Days 0 and 8 after full differentiation. Scale bars, 50μm. Error bars show means + standard deviations, with *P<0.05, **P<0.01. The p-values were obtained using the Mann-Whitney test. **(B, D)** After 0, 4, 8 and 12 days of culture in white (B) or beige (D) differentiation medium, cells were used for RNA purification. RNA samples were analysed by RT-qPCR analysis for *Egr1*, *Dcun1d3*, *Ucp1*, *Lep* and *Retn* expression in control mASC-*Egr1*^+/+^ and mutant mASC-*Egr1^−/−^* cells. Transcripts are shown relative to the level of *Rplp0* and *18S* transcripts. The relative mRNA levels were calculated using the 2^^-ΔΔCt^ method, with control being normalized to 1. For cells subjected to white differentiation, experiments were performed with n≥5 independent cultures for each genotype (n=12 at Day0, n=6 at Day4, n=5 at Day8 and n=10 at Day12 for control mASC-*Egr1*^+/+^ cells; n=11 at Day0, n=6 at Day4, n=6 at Day8 and n=12 at Day12 for mutant mASC-*Egr1^−/−^* cells). For cells subjected to beige differentiation, experiments were performed with n=16 independent cultures for each timepoint and genotype. Error bars represent the mean + standard deviations. The p-values were obtained using the Mann-Withney test. Asterisks * indicate the p-values of gene expression levels in mASC-*Egr1*^+/+^ and mASC-*Egr1^−/−^* compared to the first day of gene detection *P<0.05, **P<0.01, *** P<0.001, **** P<0.0001; # indicate the p-values of gene expression levels in mASC-*Egr1*^+/+^ versus mASC-*Egr1^−/−^*, for each time point, #P<0.05, ##P<0.01, ###P<0.001, ####P<0.0001.

Twelve days after white adipogenic stimulation, 60% of control *Egr1*^+/+^ cells have acquired a white adipocyte phenotype, marked by the presence of large lipid droplets in the cytoplasm (Fig. 4A). In contrast, only 20% of mutant mASC-*Egr1^−/−^* cells exhibited lipid droplets (Fig. 4A) suggesting that *Egr1* is necessary for white adipocyte differentiation. To confirm this hypothesis, we quantified the expression of several white and beige adipocyte markers in control and mutant cells at different days after differentiation. As expected, *Egr1* is highly expressed during the entire white adipocyte differentiation of control mASC-*Egr1*^+/+^ cells, while not detected in mutant mASC-*Egr1^−/−^* cells (Fig. 4B). The beige adipocyte marker *Ucp1* was not expressed either in control or mutant cells. During white adipocyte differentiation, *Lep*, a late marker of white adipocyte differentiation [25,26], was significantly down-regulated in mutant mASC-*Egr1^−/−^* cells compared to controls, which is consistent with the reduced white adipocyte differentiation observed in *Egr1^−/−^* cells (Fig. 4A). Resistin (*Retn*) is an adipocyte-specific hormone secreted by WAT [27]. In control cells, *Retn* expression was absent at day 0, increased during white differentiation reaching a peak at day 4, and subsequently decreased during late differentiation (Fig. 4B). A similar *Retn* expression profile has been previously described during white adipocyte differentiation of 3T3-L1 preadipocytes [28]. In contrast to their altered white differentiation phenotype (Fig. 4A), mutant mASC-*Egr1^−/−^* cells exhibited a similar *Retn* expression profile to control cells (Fig. 4B). This result indicates that *Egr1* may not regulate *Retn* expression, or that absence of *Egr1* could be compensated. In addition, it suggests that *Retn* might not be a strict marker/activator of mature white adipocyte differentiation, which has been proposed in several studies as a repressor of white adipocyte differentiation [29,30]. *Dcun1d3* displays a similar expression profile to *Retn* during white differentiation of control mASC-*Egr1*^+/+^ cells. In contrast to what we previously observed *in vivo* with the *Egr1^−/−^* mutant (Fig. 2B), *Egr1* loss-of-function did not affect *Dcun1d3* expression during mASC white adipocyte differentiation (Fig. 4B). This result can be explained by the difference between *in vivo* analysis of a mixed tissue, and study of an *in vitro* homogenous cell population strictly directed towards white differentiation.

Taken together, these results demonstrate that while mutant mASC-*Egr1^−/−^* cells express some similar markers than control cells, they lack important morphological (Fig. 4A) and transcriptomic (Fig. 4B, *Lep*) markers of normal white adipocyte differentiation. This suggests that *Egr1* has a positive role on white adipocyte differentiation.

Following beige adipogenic stimulation, control mASC-*Egr1*^+/+^ cells displayed a low level (17%) of differentiation into beige adipocytes marked by the appearance of numerous small lipid droplets in their cytoplasm (Fig. 4C). In contrast, *Egr1* deletion led to a robust differentiation of 78% of mutant mASC-*Egr1^−/−^* cells into beige adipocytes (Fig. 4C). This result shows that beige adipocyte differentiation in wild type mASC is a less efficient process compared to white differentiation (Fig. 4A, C). More importantly, this observation reveals that *Egr1* acts as a critical repressor of beige adipocyte differentiation. In wild type mASC, *Egr1* expression increases during the first time-points of beige adipocyte differentiation and decreases at the end of the process (Fig. 4C). As expected, *Ucp1* expression is significantly upregulated in mutant mACS-*Egr1^−/−^* compared to control cells, (Fig. 4D). *Lep* is not expressed during beige adipocyte differentiation of both control and mutant mASC (Fig. 4D). Expression of *Retn* is detected during beige adipocyte differentiation (Fig. 4D), while at lower levels compared to white differentiation, which further suggests that *Retn* may be a generic marker of both white and beige adipocyte differentiation [9]. Similarly, mutant mASC-Egr1^−/−^ cells which exhibit a robust differentiation rate, visualized by accumulation of lipid droplets in the cytoplasm (78%, Fig 4C), are characterized by higher *Retn* expression levels compared to wild type cells (Fig. 4D). This further confirms that *Retn* expression can be associated with beige adipocyte differentiation. *Dcun1d3* expression is increased during beige adipocyte differentiation (Fig. 4D). In addition, *Egr1* loss-of-function leads to a significant upregulation in *Dcun1d3* expression (Fig. 4D), suggesting that Egr1 represses *Dcun1d3* expression. This is similar to our *in vivo* results (Fig. 2B).

Altogether, our results indicate that *Egr1* is necessary to promote white adipocyte differentiation and to repress beige adipocyte differentiation of mASC. Consistent with the cell differentiation phenotypes, Egr1 is necessary to activate *Lep* during white differentiation and to repress *Ucp1* during beige adipocyte differentiation. *Retn* and *Dcun1d3* are expressed during both white and beige adipocyte differentiation of mASC, suggesting a generic role of these factors during adipocyte differentiation (Fig. 6). The positive role of *Egr1* on *in vitro* white adipocyte differentiation is consistent with our *in vivo* results (Figures 1, 2 and 3) and with studies correlating *Egr1* overexpression with obesity [18,19].

### *Egr1* is sufficient to promote white and to repress beige adipocyte differentiation of C3H10T1/2 mesenchymal stem cells

To further characterize the role of *Egr1* during adipocyte differentiation and confirm the results obtained with mASC, we used a second *in vitro* approach, taking advantage of the Tol2 genomic system to perform *Egr1* gain-of-function experiments in C3H10T1/2 mesenchymal cells. The Tol2 system was used in combination with the viral T2A system, which allowed stable and bicistronic expression of two proteins [31] (Fig.5-Supplement 1). Control C3H10T1/2-*Tom*-*Gfp* cells co-expressed cytoplasmic Tomato and nuclear H2B-GFP fluorescent reporter proteins, whereas in C3H10T1/2-*Tom*-*Egr1* cells, the *H2B-Gfp* coding sequence was replaced with *Egr1* coding sequence (Fig. 5-Supplement 1). C3H-*Tom*-*Gfp* and C3H-*Tom*-*Egr1* cells exhibited cytoplasmic *Tomato* expression and nuclear *H2B-Gfp* or *Egr1* expression, respectively (Fig. 5A).

**Figure 5.**
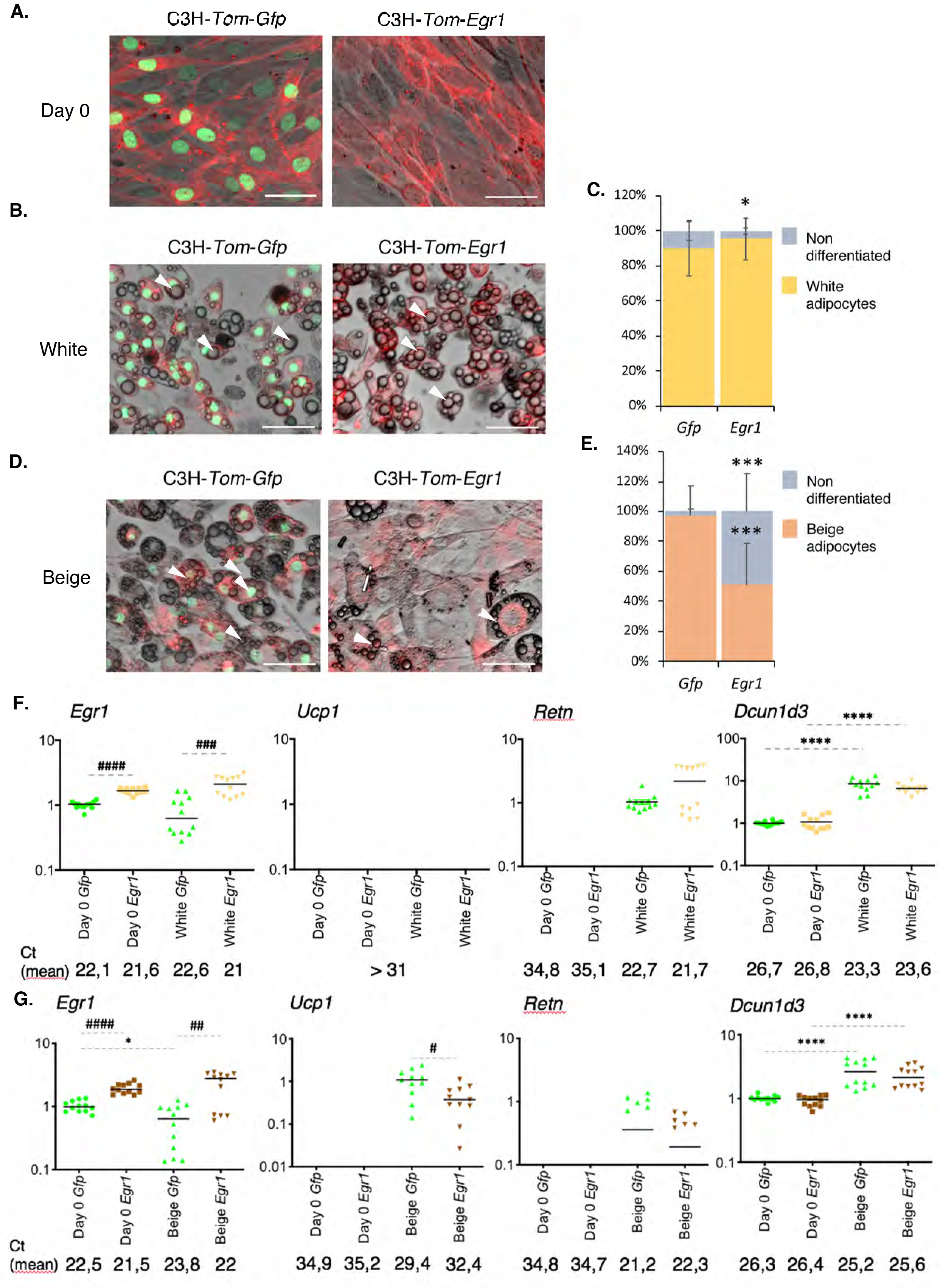
*Egr1* is sufficient to promote white adipocyte differentiation and prevent beige differentiation in a multipotent cell line. **(A-G)** Control C3H-*Tom*-*Gfp* and *Egr1*-overexpressing C3H-*Tom*-*Egr1* cells were subjected to white (B, C, F) or beige (D, E, G) differentiation for 10 and 8 days, respectively. At day 0 (A), day 10 (B, white) or day 8 (D, beige) of adipogenic culture, apotome images were obtained using brightfield, green (Gfp) and red (Tom) channels, merged, and a representative image is shown for each condition. White arrowheads indicate lipid droplets. Scale bars, 50μm. **(C, E)** C3H-*Tom*-*Gfp* and C3H-*Tom*-*Egr1* cells submitted to white (C) or beige (E) adipogenic medium were counted from n=7 (white) or n=6 (beige) pictures and expressed as a percentage of total cells differentiated or not for each condition or as total number of cells per arbitrary unit area. Error bars represent the means + standard deviations, and p-values were obtained using the Mann-Whitney test comparing differentiated and non-differentiated C3H-*Tom*-*Gfp* and C3H-*Tom-Egr1* cells, *P<0.05, ***P<0.001. **(F, G)** At Day 0 or at the end of white (F) or beige (G) differentiation, cells were used for RNA purification. RNA samples were analysed by RT-qPCR analysis for *Egr1*, *Dcun1d3*, *Ucp1* and *Retn* expression in control C3H-*Tom*-*Gfp* and *Egr1*-overexpressing C3H-*Tom*-*Egr1* cells. Transcripts are shown relative to the level of *18S* transcripts. The relative mRNA levels were calculated using the 2^^-ΔΔCt^ method, with control being normalized to 1. Experiments were performed with n=12 independent cultures for C3H-*Tom*-*Gfp* and C3H-*Tom*-*Egr1* cells for both white and beige differentiation. Error bars represent the means + standard deviations. * indicate the p-values of gene expression levels in C3H-*Tom*-*Gfp* or C3H-*Tom*-*Egr1* at the first day of gene detection versus differentiation, with*P<0.05, **** P<0.0001; # indicate the p-values of gene expression levels in C3H-*Tom*-*Gfp* versus C3H-*Tom*-*Egr1* for each time point, with #P<0.05, ##P<0.01, ###P<0.001, ####P<0.0001.

**Figure 6.**
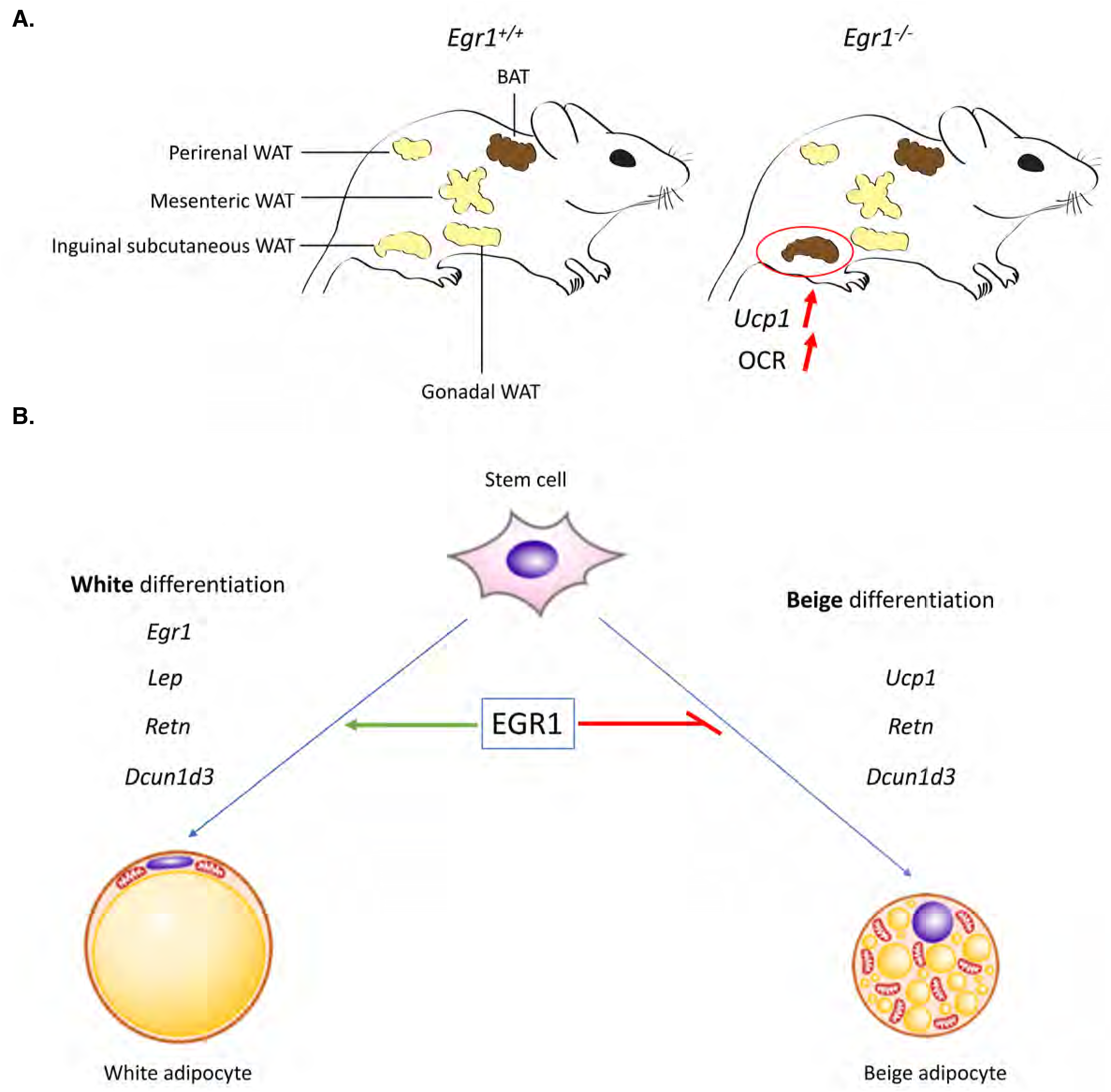
Schematic representation of the consequences of *Egr1* loss or gain-of-function on adipose tissue metabolism and adipocyte differentiation. **(A)** *Egr1* depletion leads to spontaneous and specific browning of inguinal subcutaneous white adipose tissue (WAT), visualized by increased *Ucp1* expression and oxygen consumption rate (OCR). **(B)** *Egr1* is necessary and sufficient to promote white adipocyte differentiation and reduce beige differentiation through the regulation of *Egr1*, *Ucp1*, *Lep*, *Retn* and *Dcun1d3* expression.

Ten days after white adipogenic stimulation, 90% of C3H-*Tom*-*Gfp* and 95% of C3H-*Tom*-*Egr1* cells differentiated into white adipocytes, marked by the appearance of large lipid droplets in the cytoplasm (Fig. 5B, C, white arrowheads). These observations indicate that *Egr1* gain-of-function promotes C3H10T1/2 cells differentiation into white adipocytes and support previous findings showing increased *Egr1* expression in WAT of obese mice and humans compared to lean individuals [18,19]. In contrast, while 97% of C3H-*Tom*-*Gfp* differentiated into beige adipocytes, indicated by the presence of small lipid droplets (Fig. 5D, E, white arrowheads), only 50% of C3H-*Tom*-*Egr1* cells achieved beige differentiation. These results confirm that *Egr1* gain-of-function in C3H10T1/2 mesenchymal stem cells reduces beige adipocyte differentiation [16].

To further characterize the consequences of *Egr1* gain of function during differentiation, we quantified the expression of several white and beige adipocyte markers in C3H-*Tom*-*Gfp* and C3H-*Tom*-*Egr1* cells. As expected, C3H-*Tom*-*Egr1* cells display stable overexpression of *Egr1* during adipocyte differentiation, compared to control C3H-*Tom*-*Gfp* cells (Fig. 5F, G). Similarly to our results using wild type mASC (Fig. 4D, F), control C3H-*Tom*-*Gfp* cells expressed *Egr1* throughout the white adipocyte differentiation process (Fig. 5F). Control C3H-*Tom*-*Gfp* cells also displayed a similar *Egr1* expression profile to mASC during beige adipocyte differentiation with *Egr1* downregulation being observed at the end of differentiation (Fig. 5G). This observation is consistent with the negative role of *Egr1* during beige differentiation (Fig. 4C, D; Milet *et al.*, 2017). As expected, *Ucp1* expression was highly upregulated in control C3H-*Tom*-*Gfp* cells upon differentiation into beige adipocytes (Fig. 5G). A moderate increase in *Ucp1* expression was also observed in C3H-*Tom*-*Egr1* cells when compared to control cells (Fig. 5G). Both *Retn* and *Dcun1d3* expression were upregulated in C3H-*Tom*-*Gfp* cells after white and beige differentiation (Fig. 5F, G), which strengthens the hypothesis of a generic role of these two genes during adipocyte differentiation. In addition, *Dcun1d3* expression was not altered in C3H-*Tom*-*Egr1* cells compared to controls (Fig. 5F, G), showing that *Egr1* is not required to regulate *Dcun1d3* expression during adipocyte differentiation. We could not detect any expression of the white adipocyte marker *Lep* either in C3H-*Tom*-*Gfp* or in C3H-*Tom*-*Egr1* cells during white adipocyte differentiation. This result highlights the limitations of *in vitro* cell lines in the analysis of white adipocyte differentiation and confirms that primary mASC are the only *in vitro* system that allows *Lep* detection [32].

Collectively, these results indicate that *Egr1* is sufficient to promote white adipocyte differentiation of C3H10T1/2 cells and to repress beige differentiation, presumably by downregulation of *Ucp1* expression.

In summary, *Egr1* deletion induces spontaneous WAT browning specifically in the inguinal subcutaneous depot. This process acts through *Ucp1* upregulation and results in OCR increase at a level close to the BAT (Fig. 6A). *In vitro* differentiation of mASC isolated from *Egr1* mutant SC-WAT confirmed that *Egr1* loss-of-function promotes beige adipocyte differentiation through *Ucp1* upregulation, and repress white adipocyte differentiation. We confirmed these results using *Egr1* gain-of-function experiments in C3H10T1/2 mesenchymal stem cells. Altogether, our study reveals that *Egr1* is necessary and sufficient to promote white adipocyte differentiation and to prevent beige adipocyte differentiation. These results are in accordance with the increased expression of *Egr1* observed in obese mice and humans [18,19] and establishes *Egr1* as a putative promising target to counteract obesity.

## Methods

### Mouse line

All experimental procedures using mice were conducted in accordance with the European guidelines (2010/63/UE) and were approved by the French National Ethic Committee for animal experimentation N°05 and are registered under the number 01789.02.

*Egr1* gene was inactivated in C57BL/6J mice by insertion of the *LacZ* coding sequence within the *Egr1* 5’ untranslated region, in addition to a frameshift mutation upstream of the DNA-binding domain of *Egr1* [33]. The line was maintained on a C57BL/6J genetic background (Janvier, France). Animals were bred under controlled photo-period (lights on 08:00–20:00 hours), chow diet and tap water *ad libitum*.

### RNA extraction from fat pads

Fresh inguinal subcutaneous (WAT), brown (BAT), gonadic (GAT), mesenteric (MAT) and perirenal (PAT) fat pads were removed from 8-month-old euthanized *Egr1*^+/+^ and *Egr1^−/−^* female mice and homogenized using a mechanical disruption device (Lysing Matrix A, Fast Prep MP1, 4 × 30 s, 6 m.s^−1^). Total RNA was isolated using the RNeasy mini kit (Qiagen), according to the manufacturer’s protocol, with 15 min of DNase I (Qiagen) treatment.

### Quantification of mitochondrial DNA

WAT and BAT were dissected from 8-months-old control *Egr1*^+/+^ and mutant *Egr1^−/−^* female mice and immediately frozen at −80°C. DNA purification was performed using the QIAmp DNA Minikit (Qiagen), according to the manufacturer’s instructions. DNA concentration and purity were measured using a Nanodrop. Quantitative real time PCR was performed on a StepOnePlus PCR machine (Applied Biosystems) using the FastStart Universal SYBR Green Master (Roche) with primers that amplify the mitochondrial *CytB* gene and the nuclear encoded *NDUFV1* gene as endogenous reference.

### Metabolic and bio-energetic analysis

Tissue bioenergetics were analyzed on a Seahorse extracellular flux analyzer (XF^e^24) according to the manufacturer’s instructions. 2 mg explants of WAT and BAT freshly dissected from 8-months-old control *Egr1*^+/+^ and mutant *Egr1^−/−^* female mice were immediately placed in a Seahorse XF^e^24 islet plate in 500 μL of pre-heated minimal assay medium (Seahorse Biosciences, 37 °C, pH 7.40) supplemented with glucose (2.25g/L), sodium pyruvate (1 mM) and L-glutamine (2 mM). The tissue weight was based on initial assays that optimized the oxygen consumption rate (OCR). After one hour of equilibration at 37 °C in a non-CO2 incubator, OCR was measured successively in control condition (basal rates) and after successive injections of oligomycin (1 μM) that inhibits ATP synthase (ATP-linked OCR), FCCP (10 μM) that uncouples mitochondrial OXPHOS (maximal OCR) and mixed rotenone/antimycin (10 μM) that inhibit complexes I and III (non-mitochondrial OCR).

### Detection and quantification of carbonylated proteins

WAT and BAT were dissected from 8-months-old control *Egr1*^+/+^ and mutant *Egr1^−/−^* female mice and immediately frozen at −80°C. The day of extraction, tissues were homogenized in lysis buffer (8M urea, 2M thiourea, 4% CHAPS and 10mM DTT). After 30min on ice, lysates were clarified by centrifugation at 20 000*g* for 15 min at 4°C. Protein concentrations of the supernatants were determined by the Bradford method. Labeling of protein carbonyl groups was obtained by overnight incubation of 20μg protein extracts with CF555DI hydrazide (Biotium, Fremont, CA, USA, #92165) at 4°C under medium agitation. Labeled samples were then resolved by SDS-PAGE in an anykD pre-cast gel (Biorad). After three washes using a solution 7% acetic acid and 10% ethanol, CF555DI hydrazide labeling was detected using a ChemiDoc MP Imagin System (Biorad). Total protein staining was performed using InstantBlue® Protein Stain (Expedeon). Raw data were analyzed using the Fiji/ImageJ software. In both CF555DI and Coomassie’s analysis, the protein profiles were quantified by measuring the intensity of the lane for each sample. Data were normalized using the CF555DI/Coomassie ratio.

### Isolation of murine Adipose Stem/Stromal cells (mASC)

mASC were isolated as previously described [34]. Briefly, subcutaneous inguinal fat pads were excised from wild type or *Egr1^−/−^* female mice, minced in prewarmed Krebs-ringer bicarbonate KRB (118 mM NaCl, 5mM KCL, 2.5mM CaCl2, 2mM KH2PO4, 2mM MgSO4, 25 mM NaHCO3, 5mM glucose, pH=7.4) and digested in collagenase solution (0.1%). After centrifugation, the stromal vascular fraction (SVF) was rinsed in PBS, resuspended in prewarmed Stromal medium (Dulbecco’s Modified Eagle’s Medium (DMEM; Gibco) supplemented with 10% FBS, 1% penicillin-streptomycin) and seeded on plastic flasks to allow mASC adhesion. Cells were cultured at 37 °C in humidified atmosphere with 5% CO2.

### Plasmid cloning

The *Egr1* coding sequence was obtained by PCR amplification of the plasmid *pCAß-Egr1* [35] using the following primers: m*Egr1*-Bstbl-Forward-CTAATGTTCGAAATGGCAGCGGCCAAGGC, m*Egr1*-Pmll-Reverse-ATCGTCACGTGTATTAGCAAATTTCAAT, and cloned in the transposable *pT2AL-CMV-Tom-T2A-Gfp* plasmid [31]. The *H2B-Gfp* coding sequence was removed from *pT2AL-CMV-Tom-T2A-Gfp* by BstbI and PmlI digestion and replaced with *Egr1* coding sequence,

### Transfection of C3H10T1/2 cells and isolation of *C3H-Tom-Gfp and* C3H-*Tom-Egr1* cells

Mouse mesenchymal stem cells C3H10T1/2 [36] were seeded to 6-well plates at a density 20.000 cells/cm2 in stromal medium (Dulbecco’s Modified Eagle’s Medium (DMEM; Gibco) supplemented with 10% FBS, 1% penicillin-streptomycin). Cells were transfected at 80% confluence by lipofectamine P3000 (Invitrogen) with 1μg of plasmid encoding the transposase (pT2TP) and 2μg of *pT2AL-CMV-Tom-T2A-Gfp,* or *pT2AL-CMV-Tom-T2A-Egr1,* in order to obtain stable C3H-*Tom*-*Gfp* and *C3H-Tom-Egr1* cells that overexpress *GFP* and *Egr1,* respectively. Two days after transfection, C3H-*Tom*-*Gfp* and C3H-*Tom*-*Egr1* cells were resuspended into Fluorescence Analysis Cell sorting (FACS) suspension medium (Sterile PBS 1X, 10% FBS, 1% P/S, 2mM EDTA and DAPI) and purified on a FACS Aria 3 (Becton Dickinson, CA). Live cells were gated based on the morphology, doublets were excluded and cells were sorted based on their Tomato expression.

### Cell cultures

Mouse adipose stem/stromal cells mASC-*Egr1*^+/+^ and mASC-*Egr1^−/−^*, or mouse mesenchymal stem cells C3H-*Tom*-*Gfp* and C3H-*Tom*-*Egr1* were plated on 6-well plates at a density of 33.000 cells/well in Dulbecco’s Modified Eagle’s Medium (DMEM, Invitrogen) supplemented with 10% foetal bovine serum (FBS, Sigma), 1% penicillin-streptomycin (Sigma), 1% Glutamine (Sigma), 800 μg/ml G418 Geneticin (Sigma) and incubated at 37 °C in humidified atmosphere with 5% CO2.

Cells were cultured in white adipocyte differentiation medium (STEMPRO, Thermofisher) for 12 days. Day 0 corresponds to the addition of white differentiation medium, at 90% confluence. mASC-*Egr1*^+/+^ and mASC-*Egr1^−/−^* subjected to white adipocyte differentiation medium were stopped at Day 0, Day 4, Day 8 and Day 10 or 12 for gene expression analysis and stopped at Day 10 for Oil Red O staining (as described in Milet et al., 2017). C3H-*Tom*-*Gfp* and C3H-*Tom*-*Egr1* cells subjected to white adipocyte differentiation medium were stopped at Day 0 and Day 10 or 12 for gene expression analysis.

Confluent cells were cultured in beige differentiation induction medium for 2 days and in beige maturation medium for 8 days according to published protocols [16,37]. Day 0 corresponds to the addition of beige differentiation induction medium at confluence. The maturation medium was changed every 3-4 days. mASC-*Egr1*^+/+^ and mASC-*Egr1^−/−^* subjected to beige adipocyte differentiation medium were fixed for Oil Red O staining at Day 10 or lysed for gene expression analysis at Day 0, Day 4, Day 8 and Day 10.

C3H-*Tom*-*Gfp* and C3H-*Tom*-*Egr1* cells subjected to beige adipocyte differentiation medium were stopped at Day 0 and Day 10 for gene expression analysis.

Cell number measurements were performed using the free software Fiji [38].

### Reverse-Transcription and quantitative real time PCR

Total RNAs extracted from mouse fat pads, mASC-*Egr1*^+/+^ and mASC-*Egr1^−/−^*, or C3H-*Tom*-*Gfp* and C3H-*Tom*-*Egr1* cells were Reverse Transcribed using the High Capacity Retro-transcription kit (Applied Biosystems). Quantitative PCR analyses were performed using SYBR Green PCR Master Mix (Applied Biosystems) and primers listed in Supplementary Table 1. The relative mRNA levels were calculated using the 2^^-ΔΔCt^ method [39]. The Cts were obtained from Ct normalized to *Rn18S* levels in each sample. For mRNA level analysis in fat pads, 7 to 9 independent RNA samples of 8-month-old *Egr1*^+/+^ and *Egr1^−/−^* female mice were analysed in duplicate. For mRNA level analysis of mASC-*Egr1*^+/+^ and mASC-*Egr1^−/−^* cultures, 5 to 12 independent RNA samples were analysed in duplicate for each time point. For mRNA level analysis of C3H10T1/2-*Tom*-*Gfp* and C3H10T1/2-*Tom*-*Egr1* cell cultures, 12 independent RNA samples were analysed in duplicate for each time point.

### Statistical analyses

Data was analysed using the non-parametric Mann-Withney test with Graphpad Prism V8. Results are shown as means ± standard deviations. The p-values are indicated either with the value or with * or #.

## Supporting information

Supplementary Bleher 2020

## Author contributions Statement

MB, BM, RG and YK contributed to acquisition, analysis and interpretation of data. DD contributed to analysis and interpretation of histology data and drafting the article. AL and EH contributed to conception, design, analysis and interpretation of data, drafting the article, funding.

## Acknowledgements

We thank Marthe Moldes from Centre de Recherche Saint-Antoine, Paris, France and Jacqueline Bliley, from Carnegie Mellon University, Pittsburgh, USA for comments on the manuscript. We thank Hélène Fohrer-Ting, Centre d’Histologie, d’Imagerie et de Cytométrie, Centre de Recherche des Cordeliers, INSERM, Sorbonne Université, USPC, Université Paris Descartes, Université Paris Diderot, F-75006 Paris, France, for cell sorting. We thank Alwyn Dady from Medical Sciences Institute, University of Dundee, Scotland and Sophie Gournet from IBPS, Paris, France for illustrations.

## References

1. Kershaw, E.E.; Flier, J.S. Adipose tissue as an endocrine organ. J. Clin. Endocrinol. Metab. 2004, 89, 2548–2556, doi:10.1210/jc.2004-0395.

2. Coelho, M.; Oliveira, T.; Fernandes, R. State of the art paper Biochemistry of adipose tissue: an endocrine organ. 2013, doi:10.5114/aoms.2013.33181.

3. Berry, D.C.; Stenesen, D.; Zeve, D.; Graff, J.M. The developmental origins of adipose tissue. Development 2013, 140, 3939–3949, doi:10.1242/dev.080549.

4. Sebo, Z.L.; Jeffery, E.; Holtrup, B.; Rodeheffer, M.S. A mesodermal fate map for adipose tissue. Development 2018, 145, dev166801, doi:10.1242/dev.166801.

5. Bartelt, A.; Heeren, J. Adipose tissue browning and metabolic health. Nat. Rev. Endocrinol. 2014, 10, 24–36, doi:10.1038/nrendo.2013.204.

6. Klaus S, Casteilla L, Bouillaud F, R.D. The uncoupling protein UCP: a membraneous mitochondrial ion carrier exclusively expressed in brown adipose tissue. Int J Biochem. 1991, 23, 1773883.

7. Seale, P.; Conroe, H.M.; Estall, J.; Kajimura, S.; Frontini, A.; Ishibashi, J.; Cohen, P.; Cinti, S.; Spiegelman, B.M. Prdm16 determines the thermogenic program of subcutaneous white adipose tissue in mice. 2011, 121, 53–56, doi:10.1172/JCI44271.96.

8. De Jesus, L.A.; Carvalho, S.D.; Ribeiro, M.O.; Schneider, M.; Kim, S.W.; Harney, J.W.; Larsen, P.R.; Bianco, A.C. The type 2 iodothyronine deiodinase is essential for adaptive thermogenesis in brown adipose tissue. J. Clin. Invest. 2001, 108, 1379–1385, doi:10.1172/JCI200113803.

9. Lizcano, F. The beige adipocyte as a therapy for metabolic diseases. Int. J. Mol. Sci. 2019, 20, doi:10.3390/ijms20205058.

10. Rosenwald, M.; Perdikari, A.; Rülicke, T.; Wolfrum, C. Bi-directional interconversion of brite and white adipocytes. Nat. Cell Biol. 2013, 15, 659–667, doi:10.1038/ncb2740.

11. Garcia, R.A.; Roemmich, J.N.; Claycombe, K.J. Evaluation of markers of beige adipocytes in white adipose tissue of the mouse. Nutr. Metab. (Lond). 2016, 13, 24, doi:10.1186/s12986-016-0081-2.

12. Elabd, C.; Chiellini, C.; Carmona, M.; Galitzky, J.; Cochet, O.; Petersen, R.; Pénicaud, L.; Kristiansen, K.; Bouloumié, A.; Casteilla, L.; et al. Human multipotent adipose-derived stem cells differentiate into functional brown adipocytes. Stem Cells 2009, 27, 2753–2760, doi:10.1002/stem.200.

13. Horwitz, E.M.; Le Blanc, K.; Dominici, M.; Mueller, I.; Slaper-Cortenbach, I.; Marini, F.C.; Deans, R.J.; Krause, D.S.; Keating, A. Clarification of the nomenclature for MSC: The International Society for Cellular Therapy position statement. Cytotherapy 2005, 7, 393–395, doi:10.1080/14653240500319234.

14. Zuk, P. Adipose-Derived Stem Cells in Tissue Regeneration: A Review. Int. Sch. Res. Not. 2013, 2013, e713959, doi:10.1155/2013/713959.

15. Zuk, P.A.; Zhu, M.; Ashjian, P.; De Ugarte, D.A.; Huang, J.I.; Mizuno, H.; Alfonso, Z.C.; Fraser, J.K.; Benhaim, P.; Hedrick, M.H. Human Adipose Tissue Is a Source of Multipotent Stem Cells. Mol. Biol. Cell 2002, 13, 4279–4295, doi:10.1091/mbc.E02.

16. Milet, C.; Bléher, M.; Allbright, K.; Orgeur, M.; Coulpier, F.; Duprez, D.; Havis, E. Egr1 deficiency induces browning of inguinal subcutaneous white adipose tissue in mice. Sci. Rep. 2017, 7, doi:10.1038/s41598-017-16543-7.

17. Havis, E.; Duprez, D. EGR1 Transcription Factor is a Multifaceted Regulator of Matrix Production in Tendons and Other Connective Tissues. Int. J. Mol. Sci. 2020, 21, 1–25, doi:10.3390/ijms21051664.

18. Yu, X.; Shen, N.; Zhang, M.-L.; Pan, F.-Y.; Wang, C.; Jia, W.-P.; Liu, C.; Gao, Q.; Gao, X.; Xue, B.; et al. Egr-1 decreases adipocyte insulin sensitivity by tilting PI3K/Akt and MAPK signal balance in mice. EMBO J. 2011, 30, 3754–3765, doi:10.1038/emboj.2011.277.

19. Zhang, J.; Zhang, Y.; Sun, T.; Guo, F.; Huang, S.; Chandalia, M.; Abate, N.; Fan, D.; Xin, H.-B.; Chen, Y.E.; et al. Dietary obesity-induced Egr-1 in adipocytes facilitates energy storage via suppression of FOXC2. Sci. Rep. 2013, 3, 1476, doi:10.1038/srep01476.

20. Shen, N.; Yu, X.; Pan, F.Y.; Gao, X.; Xue, B.; Li, C.J. An early response transcription factor, Egr-1, enhances insulin resistance in type 2 diabetes with chronic hyperinsulinism. J. Biol. Chem. 2011, 286, 14508–14515, doi:10.1074/jbc.M110.190165.

21. Kamohara, S.; Burcelin, R.; Halaas, J.L.; Friedman, J.M.; Charron, M.J. Acute stimulation of glucose metabolism in mice by leptin treatment. Nature 1997, 389, 374–377, doi:10.1038/38717.

22. Tartaglia, L.; M, D.; X, W.; N, D.; J, C.; R, D.; GJ, R.; LA, C.; FT, C.; J, D.; et al. Identification and expression cloning of a leptin receptor, OB-R. Cell 1995, 83, 1263–1271.

23. Trayhurn, P. Leptin--a critical body weight signal and a “master” hormone? Sci. STKE 2003, 2003, pe7, doi:10.1126/stke.2003.169.pe7.

24. Mohtar, O.; Ozdemir, C.; Roy, D.; Shantaram, D.; Emili, A.; Kandror, K. V. Egr1 mediates the effect of insulin on leptin transcription in adipocytes. J. Biol. Chem. 2019, 294, 5784–5789, doi:10.1074/jbc.AC119.007855.

25. Macdougald, O.A.; Hwang, C.S.; Fan, H.; Lane, M.D. Regulated expression of the obese gene product (leptin) in white adipose tissue and 3T3-L1 adipocytes. Proc. Natl. Acad. Sci. U. S. A. 1995, 92, 9034–9037, doi:10.1073/pnas.92.20.9034.

26. Palhinha, L.; Liechocki, S.; Hottz, E.D.; Pereira, J.A. da S.; de Almeida, C.J.; Moraes-Vieira, P.M.M.; Bozza, P.T.; Maya-Monteiro, C.M. Leptin Induces Proadipogenic and Proinflammatory Signaling in Adipocytes. Front. Endocrinol. (Lausanne). 2019, 10, doi:10.3389/fendo.2019.00841.

27. Steppan, C.M.; Bailey, S.T.; Bhat, S.; Brown, E.J.; Banerjee, R.R.; Wright, C.M.; Patel, H.R.; Ahima, R.S.; Lazar, M.A. The hormone resistin links obesity to diabetes. Nature 2001, 409, 307–312, doi:10.1038/35053000.

28. Ikeda, Y.; Tsuchiya, H.; Hama, S.; Kajimoto, K.; Kogure, K. Resistin affects lipid metabolism during adipocyte maturation of 3T3-L1 cells. FEBS J. 2013, 280, 5884–5895, doi:10.1111/febs.12514.

29. Daquinag, A.C.; Zhang, Y.; Amaya-Manzanares, F.; Simmons, P.J.; Kolonin, M.G. An Isoform of Decorin Is a Resistin Receptor on the Surface of Adipose Progenitor Cells. Cell Stem Cell 2011, 9, 74–86, doi:10.1016/j.stem.2011.05.017.

30. Kim, K.H.; Lee, K.; Moon, Y.S.; Sul, H.S. A Cysteine-rich Adipose Tissue-specific Secretory Factor Inhibits Adipocyte Differentiation. J. Biol. Chem. 2001, 276, 11252–11256, doi:10.1074/jbc.C100028200.

31. Bourgeois, A.; Esteves De Lima, J.; Charvet, B.; Kawakami, K.; Stricker, S.; Duprez, D. Stable and bicistronic expression of two genes in somite- and lateral plate-derived tissues to study chick limb development. BMC Dev. Biol. 2015, 15, 1–12, doi:10.1186/s12861-015-0088-3.

32. Wrann, C.D.; Rosen, E.D. New insights into adipocyte-specific leptin gene expression. Adipocyte 2012, 1, 168–172, doi:10.4161/adip.20574.

33. Topilko, P.; Schneider-Maunoury, S.; Levi, G.; Trembleau, a; Gourdji, D.; Driancourt, M. a; Rao, C. V; Charnay, P. Multiple pituitary and ovarian defects in Krox-24 (NGFI-A, Egr-1)-targeted mice. Mol. Endocrinol. 1998, 12, 107–122, doi:10.1210/me.12.1.107.

34. Yu, G.; Wu, X.; Kilroy, G.; Halvorsen, Y.-D.C.; Gimble, J.M.; Floyd, Z.E. Isolation of Murine Adipose-Derived Stem Cells. In Adipose-Derived Stem Cells: Methods and Protocols; Gimble, J.M., Bunnell, B.A., Eds.; Humana Press: Totowa, NJ, 2011; pp. 29–36 ISBN 978-1-61737-960-4.

35. Lejard, V.; Blais, F.; Guerquin, M.-J.; Bonnet, A.; Bonnin, M.-A.; Havis, E.; Malbouyres, M.; Bidaud, C.B.; Maro, G.; Gilardi-Hebenstreit, P.; et al. EGR1 and EGR2 involvement in vertebrate tendon differentiation. J. Biol. Chem. 2011, 286, doi:10.1074/jbc.M110.153106.

36. Reznikoff, C. a; Brankow, D.W.; Heidelberger, C. Establishment and Characterization of a Cloned Line of C3H Mouse Embryo Cells Sensitive to Postconfluence Inhibition of Division Establishment and Characterization of a Cloned Line of C3H Mouse Embryo Cells Sensitive to Postconfluence Inhibition of. 1973, 3231–3238.

37. Lone, J.; Choi, J.H.; Kim, S.W.; Yun, J.W. ScienceDirect Curcumin induces brown fat-like phenotype in 3T3-L1 and primary white adipocytes. J. Nutr. Biochem. 2015, 1–10, doi:10.1016/j.jnutbio.2015.09.006.

38. Schindelin, J.; Arganda-Carreras, I.; Frise, E.; Kaynig, V.; Longair, M.; Pietzsch, T.; Preibisch, S.; Rueden, C.; Saalfeld, S.; Schmid, B.; et al. Fiji: an open-source platform for biological-image analysis. Nat. Methods 2012, 9, 676–682, doi:10.1038/nmeth.2019.

39. Livak, K.J.; Schmittgen, T.D. Analysis of Relative Gene Expression Data Using Real-Time Quantitative PCR and the 2-ΔΔCT Method. Methods 2001, 25, 402–408, doi:10.1006/meth.2001.1262.

40. Bonnin, M.A.; Laclef, C.; Blaise, R.; Eloy-Trinquet, S.; Relaix, F.; Maire, P.; Duprez, D. Six1 is not involved in limb tendon development, but is expressed in limb connective tissue under Shh regulation. Mech. Dev. 2005, 122, 573–585, doi:10.1016/j.mod.2004.11.005.

