## Supplementary Bleher 2020 for "*Egr1* loss-of-function promotes beige adipocyte differentiation and activation specifically in inguinal subcutaneous white adipose tissue"

### Methods

#### *In situ* hybridization to adipose tissue sections

The mouse *Dcun1d3* fragment was amplified by PCR (reverse 5'ggtcatcactagccatcctaac; forward 5'cagtattcagggagggactttg) from SC-WAT cDNA and cloned in the PGEM-T easy plasmid (Promega). The antisense probe was synthesized using T7 polymerase after SalI digestion. The sense probe was obtained using Sp6 polymerase after SacII digestion. Inguinal subcutaneous fat pads were isolated from 1-month-old female mice, fixed in 4% paraformaldehyde overnight and cryo-sectioned. 8 μm wax tissue sections were used for in situ hybridization as previously described [40].

### Figure legends

**Figure 2- Figure supplement 1. *Dun1d3* is expressed in SC-WAT.** SC-WAT from wild-type mice was longitudinally cryo-sectioned. 8 μm sections were hybridized with the DIG-labeled sense and antisense probes for *Dcun1d3*. Scale bars: 100 μm.

**Figure 3- Figure supplement 1. *Egr1* loss-of-function does not affect GAT metabolic activity. (A)** Gonadal adipose tissues (GAT) of 8-month-old control *Egr1*$^{+/+}$ and mutant *Egr1*$^{-/-}$ female mice were dissected and used for DNA purification. Mitochondrial (*Cyt B*) and nuclear (*Ndufv1*) genes were quantified by qPCR and histogram represents their ratio. Error bars represent the means + standard deviations with n=3 animals for each genotype, *p<0.05. **(B, C)** Mitochondrial respiration, measured by oxygen consumption rate (OCR) in basal conditions, and after sequential addition of Oligomycin, FCCP, and a mix of Rotenone/Antimycin were simultaneously recorded on GAT tissues, freshly dissected from 8-month-old control *Egr1*$^{+/+}$ and mutant *Egr1*$^{-/-}$ female mice. **(C)** Histogram represents the basal OCR (determined as the difference between OCR before oligomycin and OCR after rotenone/antimycin A), maximal OCR (difference between OCR after FCCP and OCR after rotenone/antimycin A), ATP-linked OCR (difference between OCR before and after oligomycin), and the non-mitochondrial OCR (OCR after rotenone and antimycin A treatment) calculated from data obtained in B. (E) WAT and BAT dissected from 8-month-old control *Egr1*$^{+/+}$ and mutant *Egr1*$^{-/-}$ female mice were used to determine carbonylated protein levels.

**Figure 5- Figure supplement 1. Strategy used for *Egr1* gain-of-function in C3H10T1/2 cells. (A)** Schematic representation of the T2A control vector characterized by the presence of the two reporter fluorescent genes *Tomato* and *H2B-GFP*, separated by the T2A peptide and flanked by the Tol2 genomic integration system. In this vector, the *H2B-GFP* coding sequence has been replaced by the *Egr1* coding sequence to allow *Egr1* overexpression. **(B)** Strategy used for stable and bi-cistronic expression of *Tomato* and *H2B-GFP* or *Egr1*. C3H10T1/2 cells were co-transfected with a first vector containing the transposase coding sequence and with the T2A-H2B-GFP or the T2A-Egr1 vector. Transposase expression leads to stable integration of the CMV/βactin promoter-Tomato-T2A-H2B-GFP or CMV/βactin promoter-Tomato-T2A-EGR1 transgenes into the cell genome. Expression of both Tomato-T2A-H2B-GFP and Tomato-T2A-EGR1 cassettes are under the control of the CMV/βactin promoter and leads to the transcription of one single mRNA. During translation, the self-cleavage of the T2A peptide allows the production of the two proteins Tomato and H2B-GFP or Tomato and EGR1 in stoichiometric proportions. Tomato and H2B-GFP localizations are cytoplasmic and nuclear, respectively [31].

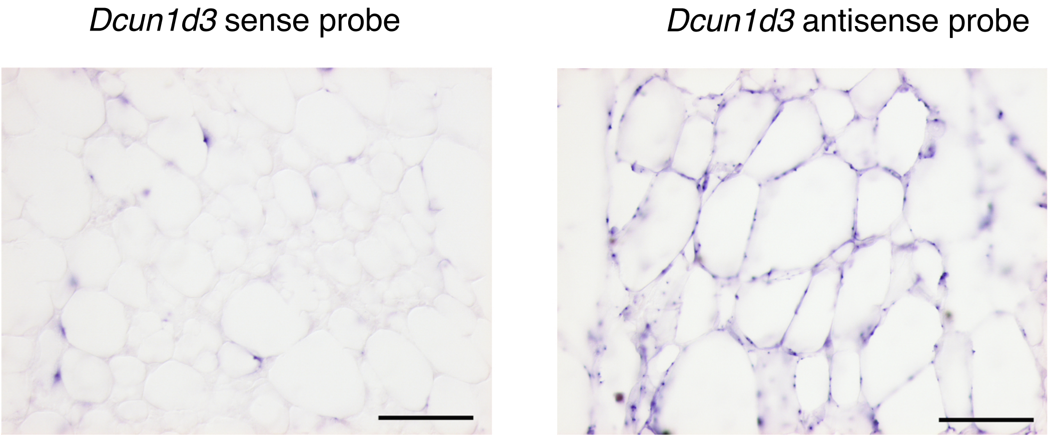


**Figure 2 -figure supplement 1**

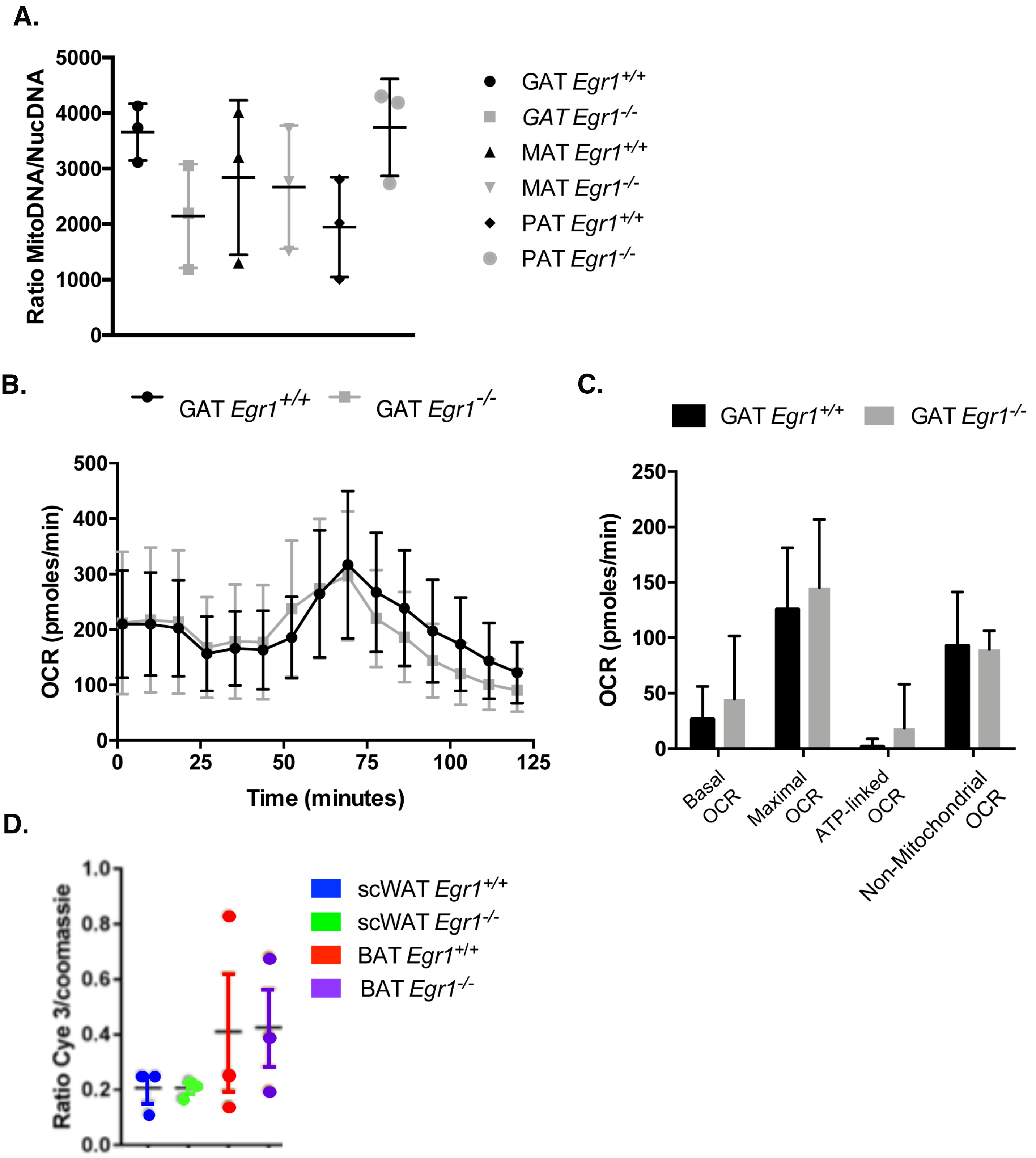


**Figure 3-figure supplement 1**

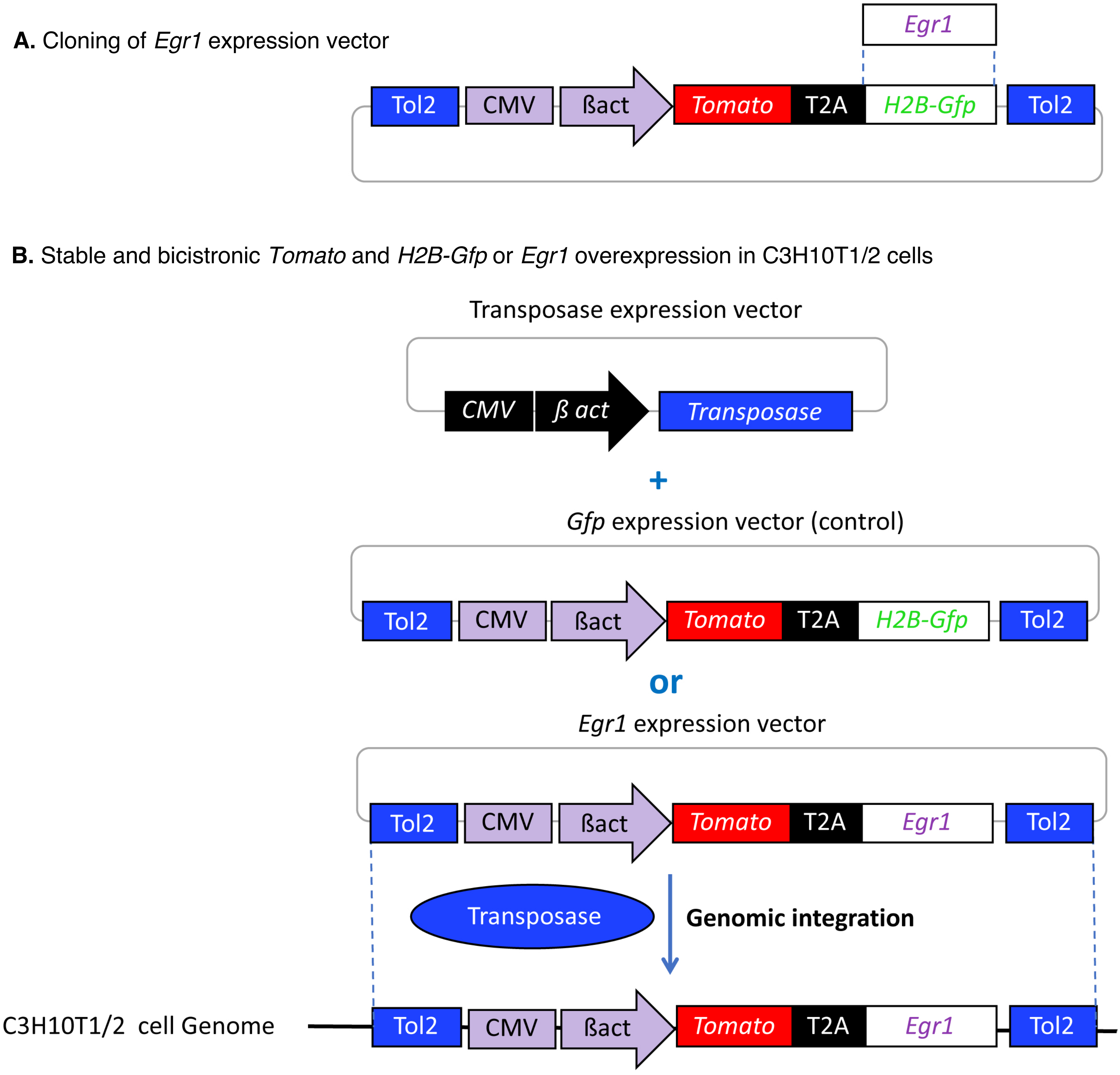


**Figure 5-figure supplement 1**

### Supplementary Table 1

| **Primers for RT-qPCR analysis** | | |
|---|---|---|
| **Gene name** | **Forward Primer** | **Reverse Primer** |
| ***Dcun1d3*** | 5’-GCTGACTCTGTCCTACCTTATTC | 5’-CCACCCACTTTCCAGTACAT |
| ***Egr1*** | 5’-CAGCGCCTTCAATCCTCAAG | 5’-GCGATGTCAGAAAAGGACTCTGT |
| ***Lep*** | 5’-TACCGCATTTCAGGGCACAT | 5’- CCCAGGTATCCCGTGTCAAC |
| ***Retn*** | 5’-GCCATCGACAAGAAGATCAA | 5’-CTTCCCTCTGGAGGAGACTG |
| ***Rn18S*** | 5’-GGCGACGACCCATTCG | 5’-ACCCGTGGTCACCATGGTA |
| ***Rplp0*** | 5’- ACCTCCTTCTTCCAGGCTTT | 5’- CTCCCACCTTGTCTCCAGTC |
| ***Ucp1*** | 5’- GGGCATTCAGAGGCAAATCAGCTT | 5’- ACACTGCCACACCTCCAGTCATTA |
